# The class I TCP transcription factor AtTCP8 is a modulator of phytohormone-responsive signaling networks

**DOI:** 10.1101/2021.12.21.473710

**Authors:** Benjamin J. Spears, Samuel A. McInturf, Meghann Chlebowski, Jianbin Su, David G. Mendoza-Cózatl, Walter Gassmann

**Affiliations:** Division of Plant Sciences, University of Missouri, Columbia, Missouri, United States of America; Christopher S. Bond Life Sciences Center and Interdisciplinary Plant Group, University of Missouri, Columbia, Missouri, United States of America; Department of Biological Sciences, Butler University, Indianapolis, Indiana, United States of America

## Abstract

The plant-specific TEOSINTE BRANCHED1/ CYCLOIDEA/ PROLIFERATING CELL FACTOR (TCP) transcription factor family is most closely associated with regulating plant developmental programs. Recently, TCPs were also shown to mediate host immune signaling, both as targets of pathogen virulence factors and regulators of plant defense genes. However, any comprehensive characterization of TCP gene targets is still lacking. Loss of the class I TCP *AtTCP8* attenuates early immune signaling, and when combined with mutations in *AtTCP14* and *AtTCP15*, additional layers of defense signaling in *Arabidopsis thaliana*. Here we focus on TCP8, the most poorly characterized of the three to date. We use chIP and RNA-sequencing to identify TCP8-bound gene promoters and differentially regulated genes in the *tcp8* mutant, data sets that are heavily enriched in signaling components for multiple phytohormone pathways, including brassinosteroids (BRs), auxin, and jasmonic acid (JA). Using BR signaling as a representative example, we show that TCP8 directly binds and activates the promoters of the key BR transcriptional regulators *BZR1* and *BZR2/BES1*. Furthermore, *tcp8* mutant seedlings exhibit altered BR-responsive growth patterns and complementary reductions in *BZR2* transcript levels, while the expressed protein demonstrates BR-responsive changes in subnuclear localization and transcriptional activity. We conclude that one explanation for the significant targeting of TCP8 alongside other TCP family members by pathogen effectors may lie in its role as a modulator of brassinosteroid and other plant hormone signaling pathways.

**One Sentence Summary:** One member of a pathogen-targeted transcription factor family modulates phytohormone response networks and displays brassinosteroid-dependent cellular location and activity.

## INTRODUCTION

Plants must be able to perceive diverse local environmental conditions and integrate that data into an appropriate biological response. In addition to abiotic stresses like light, water, and nutrient availability, a successful plant must also be able to respond to the presence of a variety of different pests and pathogens to protect itself from disease. The metabolic costs of these distinct biological processes mandate tight control to allow plants to respond appropriately to stresses without unnecessary costs to growth and development that arise from unregulated signaling (Couto and Zipfel, 2016). These balances are largely governed through a complex network of phytohormone signaling pathways (Shigenaga et al., 2017). This tradeoff is exemplified by yield loss in crop species and the model plant *Arabidopsis thaliana* (Arabidopsis) conferred through enhanced immune signaling (Ning et al., 2017). Even more striking is the severe growth inhibition observed in Arabidopsis plants continuously exposed to immune elicitors, or mutants constitutively expressing defense genes (van Wersch et al., 2016). It is likely not only the intensity of either signaling pathway, but rather the tactical precision with which they are activated that may determine the success of a plant. Small perturbations in phytohormone status can therefore dramatically influence a plant’s ability to defend itself, its growth potential, or both. This is a principle that pathogens have evolved to exploit to their benefit through the secretion of host-modulating virulence factors (Ma and Ma, 2016).

The plant immune system is a multilayered signaling network that offers robust protection against infection by most pathogens. Plants deploy a suite of pattern recognition receptors (PRRs), among them the leucine-rich repeat receptor-like kinases (LRR-RLKs) FLS2 and EFR, to the cell surface that interact with specific sets of pathogen-associated molecular patterns (PAMPs, MAMPs) (Macho and Zipfel, 2014). Upon detection of a potential extracellular threat, LRR-RLKs transmit the signal to the cell interior through their kinase domains, activating a signal transduction cascade that extends to the nucleus. There, the activities of transcriptional regulators are altered, likely through corresponding changes in protein modifications, stability, interactions, and localization, to modulate gene expression towards the production of diverse physiological responses, ultimately restricting pathogen virulence and growth. This collective response is known as PAMP-triggered immunity (PTI). Inactivation of this immune response through targeting and disabling of critical signaling components by secreted effector proteins is a major strategy by which plant pathogens may promote their own virulence (Zhou et al., 2014; Ahmed et al., 2018; Ceulemans et al., 2021).

Immune signaling has only recently been folded into the transcriptional repertoire of the 24-member *TEOSINTE BRANCHED1/ CYCLOIDEA/ PROLIFERATING CELL FACTOR* (TCP) transcription factor family in Arabidopsis, but regulation is evident at multiple levels (Kim et al., 2014; Yang et al., 2017; Li et al., 2018; Zhang et al., 2018; Spears et al., 2019). TCPs have largely been characterized as regulators of many facets of plant growth and development in a semi-redundant fashion, among others cell elongation and proliferation, stature, germination, flowering time, pollen development, and leaf morphology (Li, 2015). In support of these diverse developmental roles, TCPs have also been implicated in the direct regulation of both biosynthesis and signaling pathways of many plant hormones, including jasmonic acid (JA), salicylic acid (SA), cytokinin (CK), ABA, auxin, and brassinosteroids (BR) (Schommer et al., 2008; Guo et al., 2010; Mukhopadhyay and Tyagi, 2015; Wang et al., 2015; Gonzalez-Grandio et al., 2017). As seen with related basic helix-loop-helix (bHLH) TFs like the PHYTOCHROME INTERACTING FACTOR (PIF) and BRASSINAZOLE-RESISTANT (BZR) families, heterodimeric interactions between TCPs of either class and with other TF families hints at a complex mechanism of regulation that may function to ensure the environmentally proper composition of plant hormone signaling (Danisman et al., 2013). It is fitting, then, that several TCP family proteins have been identified as the targets of secreted effector proteins from multiple pathogens (Sugio et al., 2011; Weßling et al., 2014; Lopez et al., 2015; Yang et al., 2017; González-Fuente et al., 2020), potentially as part of an evolved strategy to promote virulence through modulation of these phytohormone signaling pathways.

Previous studies have described redundancies in regulation of host immunity by the trio of class I members TCP8, TCP14, and TCP15 (Kim et al., 2014; Li et al., 2018; Spears et al., 2019). However, recent work has pointed towards individual modes of action for each of these TCPs in the regulation of unique sets of defense and development-related phytohormone signaling components. TCP14 directly suppresses JA signaling (Yang et al., 2017) and TCP15 activates the transcription of *PR5* and *SNC1* (Li et al., 2018; Zhang et al., 2018), and TCP8 the transcription of *ICS1* (Wang et al., 2015) to promote SA signaling to the same effect. TCP15 and TCP14 are often implicated together in the control of cytokinin and GA-dependent cell division and germination (Steiner et al., 2012; Lucero et al., 2015; Gastaldi et al., 2020), but TCP8 is generally uninvolved. Of the trio, TCP8 has been relatively understudied-particularly in the context of growth and development-related functions.

Here, we aimed to further characterize the activities of TCP8 individually; to this end, we performed chromatin immunoprecipitation (chIP-seq) and RNA sequencing (RNA-seq) to identify patterns of genome-wide TCP8-bound promoters and differential gene expression between WT, *tcp8* single and *tcp8 tcp14 tcp15* triple (*t8t14t15*) mutants. We describe the enrichment of phytohormone signaling genes in multiple pathways and further validate our sequencing data by characterizing a novel role for TCP8 in contributing to the regulation of BR signaling. Our data show that TCP8 directly binds and transcriptionally activates key BR gene promoters *in planta*, that the activity and subcellular localization of TCP8 is BR-dependent, and through 24-epiBL (BL) insensitivity and brassinazole (Brz) hypersensitivity assays, that *tcp8* is compromised in BR-signaling. Additionally, we demonstrate that TCP8 interacts directly with master BR regulators BZR1 and BZR2 in multiple expression systems, findings that support the notion of TCP heterodimerization with other TF families in a complex regulatory module to broadly regulate BR and other hormone signaling pathways in Arabidopsis.

## RESULTS

### Differential regulation of hormone signaling pathways by class I TCPs

We generated a genome-wide profile of TCP8-interacting gene promoters to clarify mechanisms by which TCP8 is capable of regulating both defense and developmental signaling. ChIP-seq in a previously characterized native promoter-driven *pTCP8:TCP8-HA* [*t8t14t15*] transgenic line (Spears et al., 2019) was used to identify approximately 3500 TCP8 binding sites across the genome (Table S1). Although 16% of identified candidate peaks (568) were located upstream of the TSS of gene loci, most peaks were located in non-promoter regions (Figure 1A, Supplemental Table S2). This is not an unusual feature of some transcription factors, especially in the bHLH family (Heyndrickx et al., 2014). For the purpose of our study, we focused on the subset of genes with promoter-localized binding sites.

**Figure 1.**
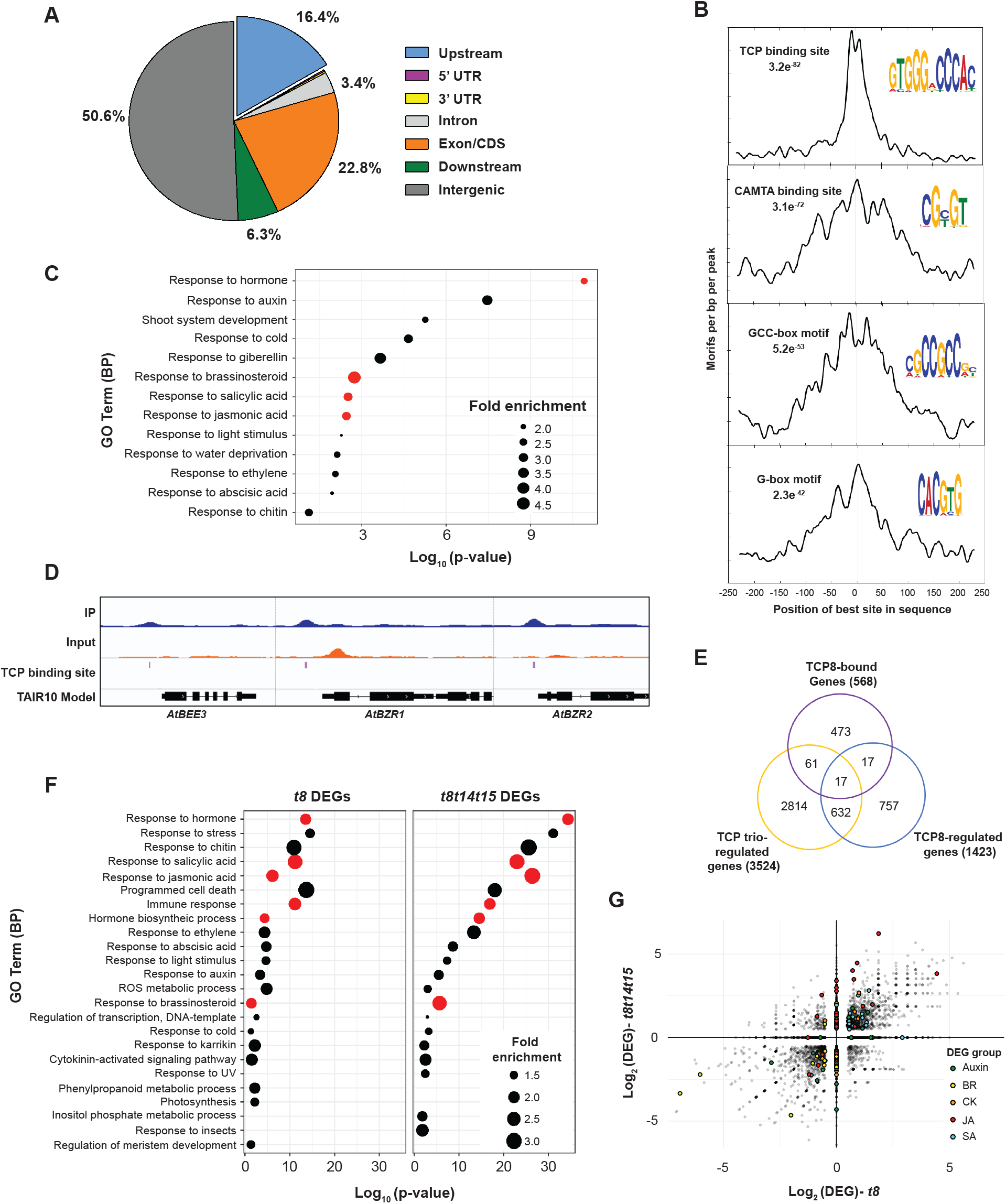
TCP8 genomic targets are overrepresented in brassinosteroid and other phytohormone signaling pathways. A, Genomic distribution of TCP8 binding peaks, with peaks up to −5 kb classified as upstream. B, Enrichment of known transcription factor binding motifs in 250 bp flanking sequences around promoter-localized TCP8 binding peaks was determined by MEME-Suite tools. Highest interesting enriched motifs are represented by sequence logos. C, Gene ontology (GO) analysis of promoter-localized TCP8-regulated gene candidates. Log10 (P-values) represented on the x-axis. Categories in red are highlighted in the text/are discussed further. D, Representative TCP8 binding peaks in notable BR gene promoters *AtBEE3, AtBZR1*, and *AtBZR2*. E, Limited overlap observed between TCP8-bound regulatory targets and *tcp8* DEGs identified by RNA-seq. Greater overlap is observed between *tcp8* and *t8t14t15* DEGs, but the majority of *tcp8* DEGs are exclusive to that genetic background. F, GO analysis of *TCP8* and *T8T14T15*-regulated genes. Log10 (P-values) are represented on the x-axis. Categories in red are highlighted in the text/are discussed further. G, Scatter plot of log2 fold change values in *tcp8*/Col-0 (x-axis) and *t8t14t15*/Col-0 (y-axis) RNA-seq DEGs. Genes associated with phytohormone groups are highlighted in green (auxin), yellow (BR), orange (CK), red (JA), or blue (SA).

From these candidate binding sites, we scanned the surrounding 500 bp regions for identification of enriched sequence motifs (Figure 1B). As expected, the most significantly enriched motif (GTGGGxCCCAC) corresponds to a canonical class I TCP binding site (O’Malley et al., 2016). Other notable motifs include the binding site for the immunity-suppressing CAMTA TF family (CGCGT) and the GCC-box (GCCGCC) of AP2/EREBP factors that regulate ethylene and ABA signaling (Dietz et al., 2010). We were particularly drawn to the G-box (CACGTG) associated with transcription factors of the PIF and BZR families, known for regulation of plant growth and response to environmental light conditions. Notably, genetic interactions between BZRs/PIFs and TCPs have been well-characterized (Perrella et al., 2018; Zhou et al., 2018; Ferrero et al., 2019; Zhou et al., 2019).

A gene ontology (GO) term enrichment analysis identified significant enrichment of genes involved in BR signaling, as well as most major phytohormone signaling pathways, including strong enrichment of auxin-related genes (Figure 1C). Given the established role of TCP8 in the host immune response, the enrichment of SA-regulatory genes such as *WRKY40* are unsurprising and highlight these candidates as potential causal factors for future study. For BR-related genes tested, peaks correlated with the presence of a TCP binding site (Figure 1D). We have highlighted the largest peak heights and most notable pathway representative candidates of these categories (Table 1).

**Table 1.** Phytohormone signaling regulators identified by chIP-sequencing as candidate gene targets of AtTCP8

| TAIR ID | Gene Description | Peak Height | TAIR ID | Gene Description | Peak Height |
| --- | --- | --- | --- | --- | --- |
| Auxin |  |  | Salicylic Acid |  |  |
| AT1G53700 | Serine/threonine-protein kinase;WAG1 | 203 | AT2G04880 | WRKY transcription factor 1; WRKY1 | 120 |
| AT5G59430 | TELOMERE REPEAT BINDING PROTEIN 1 (TRP1) | 201 | AT1G80840 | WRKY transcription factor 40; WRKY40 | 112 |
| AT5G19140 | Aluminum induced protein with YGL and LRDR motifs;AILP1 | 189 | AT4G05320 | POLYUBIQUITIN 10 (UBQ10) | 90 |
| AT3G14370 | Serine/threonine-protein kinase; WAG2 | 176 | AT2G20580 | 26S proteasome subunit 2 homolog A; RPN1A | 66 |
| AT3G04730 | Auxin-responsive protein; IAA16 | 169 | AT3G48090 | ENHANCED DISEASE SUSCEPTIBILITY 1 (EDS1); EDS1A | 58 |
| AT5G62000 | AUXIN RESPONSE FACTOR 2 (ARF2) | 159 | Jasmonic Acid |  |  |
| AT1G15750 | TOPLESS (TPL) | 150 | AT2G18790 | Phytochrome B (PHYB) | 101 |
| AT4G37610 | BTB/POZ and TAZ domain-containing protein 5; BT5 | 116 | AT3G15510 | NAC transcription factor 56; NAC056 | 176 |
| AT3G61830 | AUXIN RESPONSE FACTOR 18 (ARF18) | 110 | AT5G64810 | WRKY transcription factor 51; WRKY51 | 71 |
| AT1G04240 | Auxin-responsive protein; IAA3 | 110 | Ethylene |  |  |
| AT4G29080 | Auxin-responsive protein; IAA27 | 104 | AT1G06160 | Ethylene-responsive transcription factor 94; ERF94 | 161 |
| AT1G35240 | AUXIN RESPONSE FACTOR 20 (ARF20) | 97 | AT5G61590 | Ethylene-responsive transcription factor 107; ERF107 | 141 |
| AT2G39550 | Geranylgeranyl transferase type-1 subunit beta; GGB | 95 | AT1G62300 | WRKY transcription factor 6; WRKY6 | 140 |
| AT4G32880 | Homeobox-leucine zipper protein; ATHB-8 | 91 | AT1G64060 | Respiratory burst oxidase homolog protein F;RBOHF | 130 |
| AT4G16780 | Homeobox-leucine zipper protein; HAT4 | 89 | AT3G15210 | Ethylene-responsive transcription factor 4; ERF4 | 125 |
| AT2G42620 | F-box protein; MAX2 | 85 | AT5G38480 | 14-3-3-like protein GF14 psi; GRF3 | 118 |
| AT3G48360 | BTB/POZ and TAZ domain-containing protein 2; BT2 | 81 | AT5G25190 | Ethylene-responsive transcription factor 3; ERF3 | 98 |
| Gibberellin |  |  | AT4G23750 | Ethylene-responsive transcription factor; CRF2 | 75 |
| AT4G19700 | E3 ubiquitin-protein ligase; BOI | 178 | Abscisic Acid |  |  |
| AT1G76180 | Dehydrin; ERD14 | 123 | AT3G22380 | TIME FOR COFFEE (TIC) | 205 |
| AT1G69530 | Expansin-A1 (EXPA1) | 108 | AT1G20440 | Dehydrin; COR47 | 189 |
| AT3G63010 | Gibberellin receptor; GID1B | 98 | AT1G20450 | Dehydrin; ERD10 | 157 |
| AT3G58070 | Zinc finger protein; GIS | 90 | AT3G07360 | U-box domain-containing protein 9; PUB9 | 153 |
| AT5G11260 | LONG HYPOCOTYL 5 (HY5) | 72 | AT1G69270 | LRR receptor-like serine/threonine-protein kinase; RPK1 | 150 |
| Brassinosteroid |  |  | AT1G75240 | Zinc-finger homeodomain protein 5; ZHD5 | 132 |
| AT1G13260 | AP2/ERF and B3 domain-containing transcription factor; RAV1 | 241 | AT1G56600 | GALACTINOL SYNTHASE 2 (GOLS2) | 125 |
| AT4G28720 | Probable indole-3-pyruvate monooxygenase; YUCCA8 (YUC8) | 145 | AT2G47180 | GALACTINOL SYNTHASE 1 (GOLS1) | 123 |
| AT2G42870 | PHYTOCHROME RAPIDLY REGULATED 1 (PAR1) | 134 | AT5G67450 | Zinc finger protein AZF1;AZF1 | 98 |
| AT1G75080 | BRASSINAZOLE-RESISTANT 1 (BZR1) | 125 |  |  |  |
| AT1G19350 | BRASSINAZOLE-RESISTANT 2 (BZR2);BES1 | 124 |  |  |  |
| AT4G39400 | BRASSINOSTEROID INSENSITIVE 1 (BRI1) | 76 |  |  |  |
| AT5G08130 | BES-INTERACTING MYC-LIKE PROTEIN 1 (BIM1) | 71 |  |  |  |

To further interrogate TCP8-specific signaling roles, we performed RNA-seq analysis comparing total mRNA collected from Col-0, *tcp8*, and *t8t14t15* seedlings under our chIP-seq growth conditions. Differential expression relative to Col-0 was observed for 1423 genes in *tcp8* and 3524 genes in *t8t14t15* (Figure 1E, Table S3). In general, these mutations had relatively low magnitude of effect overall on global gene expression; this was not unexpected, given the redundancies with which this TF family functions. However, only 44% of the differentially expressed genes (DEGs) in *tcp8* were also affected in *t8t14t15*, with similar distributions seen when comparing grouped TCP-upregulated and TCP-downregulated DEGs (Supplemental Figure S1). This would support the TCP trio having sets of both distinct and common regulatory targets and roles.

Surprisingly, only 6% and 8% of candidates identified by chIP-seq as having TCP8-bound promoter regions were shown to be differentially expressed in *tcp8* and *t8t14t15*, respectively. Similar observations have previously been made for the related bHLH PIF3 (Zhang et al., 2013), which highlights the complex, likely heterodimeric and additive mechanisms of regulation by TCP8 and other TCPs. The subset of 34 genes bound by TCP8 and differentially expressed in the *tcp8* mutant did not fall into any clear functional or physiological categories but nonetheless represents a valuable resource for direct transcriptional regulatory targets to be pursued individually in future studies (Table S4). Specific environmental conditions or hormone treatments may be required to effectively capture these alterations to the transcriptional profile. For the purpose of this study, we continued to focus on more general sectors regulated by TCP8.

A GO term analysis of TCP8 and TCP8/TCP14/TCP15-regulated genes identified significant enrichment of phytohormone signaling genes associated with growth, defense, and light responses in both sets (Figure 1F). The dramatically increased enrichment of JA signaling genes in *t8t14t15* relative to *tcp8* is unsurprising as previous studies have described the suppression of JA and enhancement of SA signaling by TCP14 and TCP15. We observed a broad pattern of increased enrichment of DEGs associated with most hormones in the *t8t14t15* background relative to *tcp8*, but this trend is heavily muted for pathways like auxin, ABA, and BR, where loss of *TCP8* alone was sufficient to observe similarly weak enrichment of those DEG groups.

A comparison of absolute fold changes within a curated list of phytohormone-related DEG candidates in *tcp8* and *t8t14t15* (Figure 1G) also supports the idea of specialization between the TCPs tested. In aggregate, more significant *tcp8*-specific changes were observed for auxin-related genes, while differences in BR and SA-related genes were smaller but still significant, cytokinin (CK)-related genes were largely uncoordinated, and JA-specific genes were clearly *tcp14* and *tcp15*-specific. We decided to further probe the BR signaling pathway as a test case for candidates identified in the chIP dataset, exploring the possibility of relatively small perturbations of phytohormone-related gene expression levels leading to significant physiological consequences. BR-related DEGs stood out by predominantly being lower in *tcp* mutants and represented some of the most down-regulated DEGs. Additionally, intersections between the BR and SA response are well-established and the known regulation of SA signaling by TCP8 may uniquely position it to modulate balance of these pathway outputs.

### TCP8 binds and activates BR gene promoters *in planta*

Putative TCP8 regulatory targets are associated with perception of BR at the plasma membrane (*BRI1*), transcriptional output (*BZR1, BZR2, BIM1, BEE3*) and biosynthesis of BR (*BRX*). As class I TCPs have yet to be directly implicated in the regulation of the BR pathway, we explored several of these candidates further to confirm a similar level of regulation by a class I TCP. For a subset of BR signaling candidates, we aimed to verify the chIP-seq results through targeted chIP-qPCR amplification of regions flanking identified TCP binding sites. Clear enrichment of the target regions of *BZR1, BZR2*, and *BRI1* promoters was observed relative to a non-target region after immunoprecipitation with HA-directed antibody (Figure 2A). Non-BR genes identified in the chIP-seq dataset were also pulled down, here verified by enrichment of the *WRKY40* promoter (Figure 2B).

**Figure 2.**
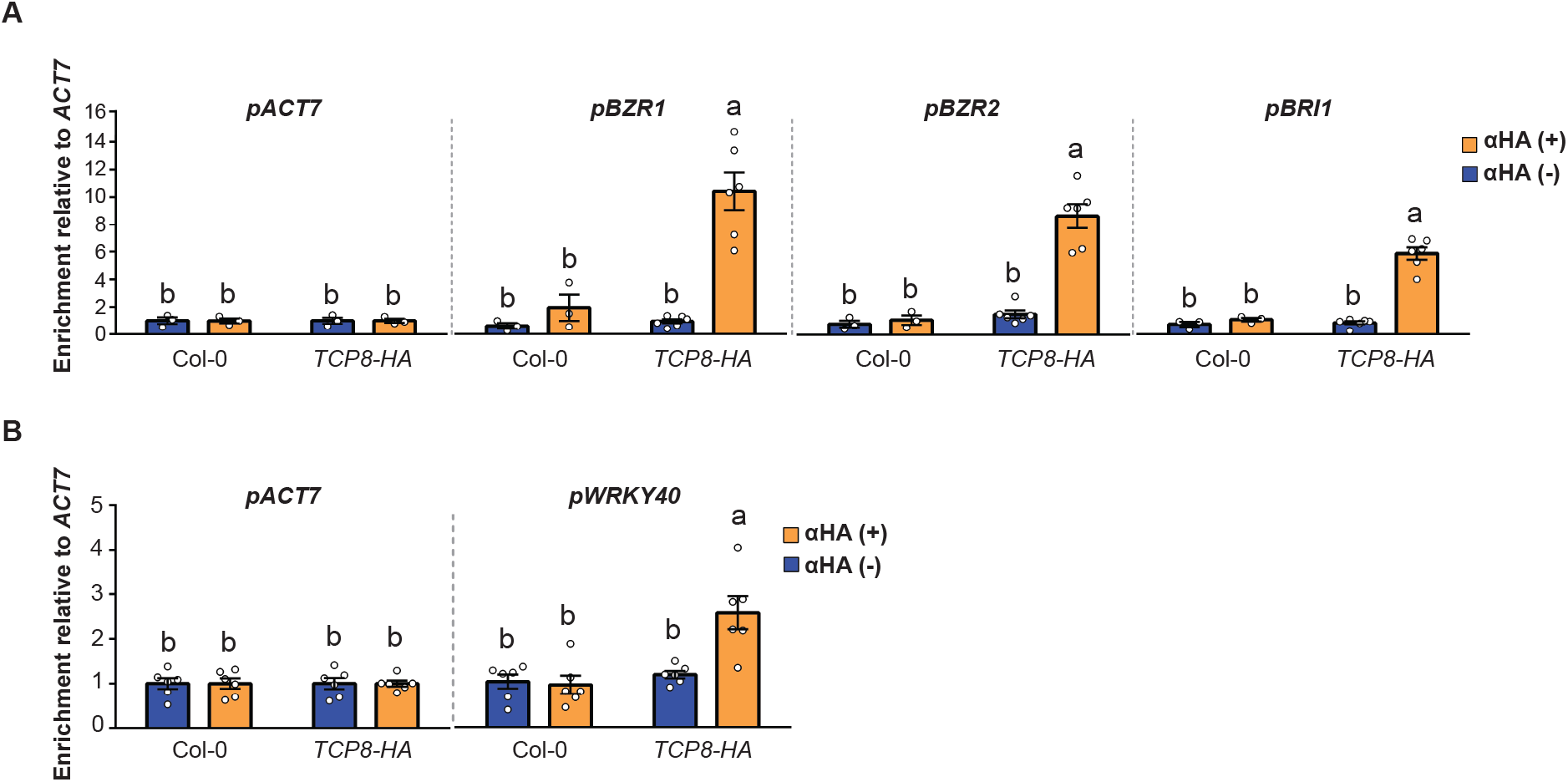
TCP8-bound BR gene regulatory targets verified by chIP-PCR. A, Chromatin of Col-0 seedlings was immunoprecipitated with either αHA (HA+) or protein A (HA-) beads and enrichment of BR genes determined relative to the non-target *ACT7* promoter (*n* = 3 Col-0, *n* = 6 *pT8:T8-HA*). B, Verification of *WRKY40* promoter occupancy by TCP8 as representative SA gene candidate. Enrichment of *WRKY40* promoter was performed in the same manner as in (A), confirming that regulatory candidates from multiple phytohormone pathways are occupied by TCP8 (*n* = 6). For all chIP-PCR experiments, error bars indicate SE, significance (ANOVA) with Tukey multiple comparison test, letters denote difference at *P* <0.05.

To determine if the direct interaction between TCP8 and BR gene promoters is sufficient for their activation, we transiently co-expressed *35S:HA-TCP8* with *promoter:GUS* reporter constructs in *N. benthamiana* leaf cells. Through quantification of GUS activity, we demonstrated that transiently expressed HA-TCP8 is capable of strongly activating *BZR1* and *BZR2* promoters *in planta* (Figure 3A,B). Mutating two identified TCP binding sites of the *BZR2* promoter strongly attenuated activation by HA-TCP8. Similar mutations in the *BZR1* promoter showed little effect on its activation, possibly due to the presence of other compensating TCP binding sites that were not interrogated in this study. These data suggest that TCP8 directly binds and activates the promoters of several key BR regulatory genes, likely as a positive regulator of BR signaling.

**Figure 3.**
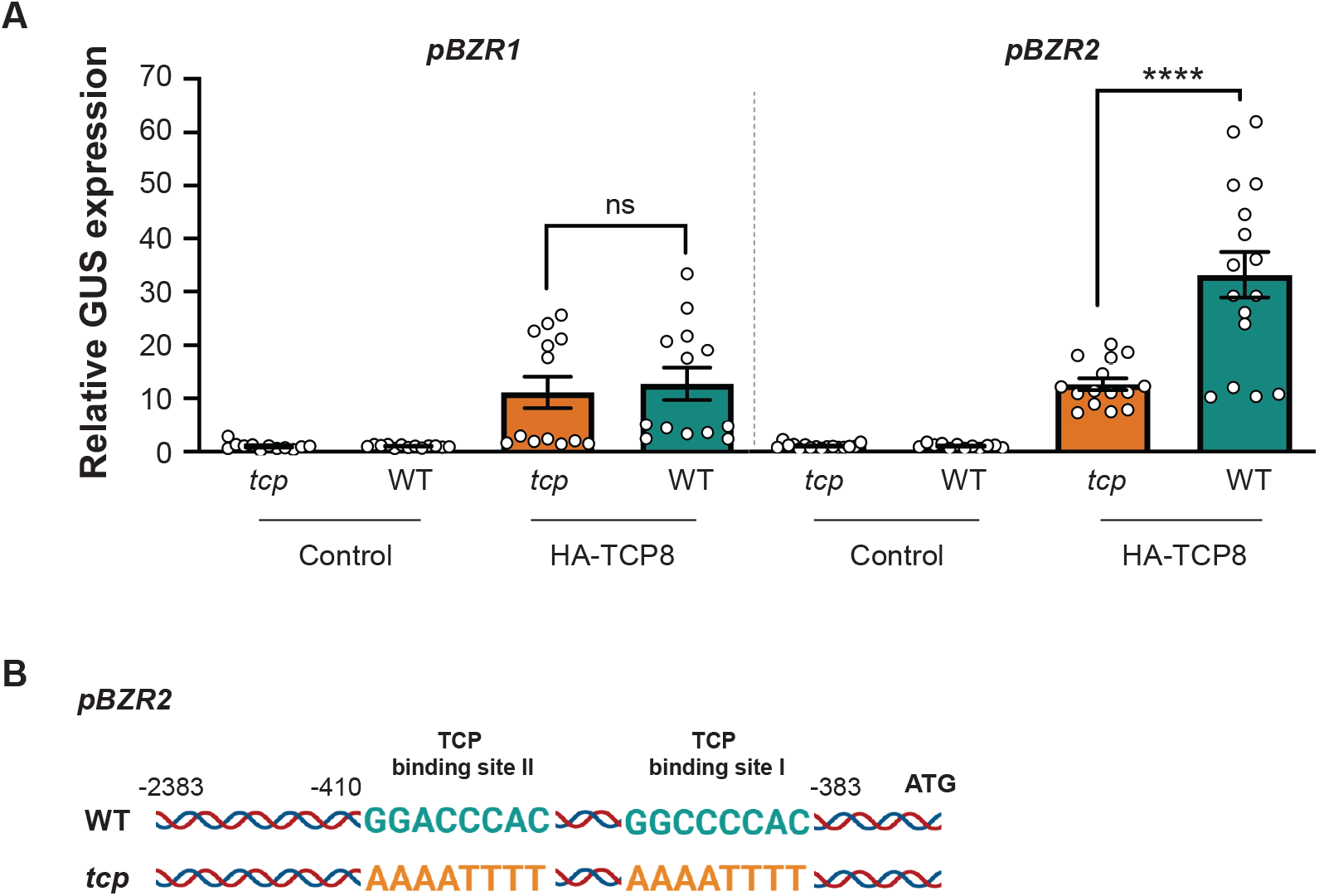
TCP8 transactivates *pBZR2* promoter:GUS construct in a TCP binding-site dependent manner in *N. benthamiana*. A, GUS expression relative to control was determined by 4-MUG assay after co-expression of reporter construct and EV (control) or HA-TCP8 construct in *N. benthamiana* cells and data combined from three independent experiments. Error bars indicate SE, *\*\*\*\*P* <0.0001 (*n* = 13-16, Student’s *t*-test). B, Model of the wild-type (WT) and TCP8-binding site mutant (*tcp*) promoter versions tested in the GUS reporter construct.

### Enhanced reactive oxygen species (ROS) production in the *tcp8* mutant is suppressed by BR treatment

In an earlier study, we observed an increase in elf18-induced ROS production in *t8t14t15*, despite impaired downstream PTI outputs like SA marker gene induction and resistance to pathogen infection (Spears et al., 2019). Notably, the tradeoff between growth and defense is demonstrated by a well-established and multileveled antagonism between BR and immune signaling, the mechanisms for which are not yet fully understood but are thought to depend on BZR activities (Belkhadir et al., 2012; Lozano-Duran et al., 2013). We therefore suspected that impaired BR signaling in *tcp8* could be a contributing factor to the enhanced ROS phenotype. In support of this hypothesis, we now find that the loss of *TCP8* alone is sufficient for enhanced ROS in a standard leaf disc assay (Figure 4A). The lack of noticeable changes in downstream (‘late’) PTI outputs in *tcp8* (Supplemental Figure S2) would suggest that suppression of early PTI signaling may reflect a *TCP8-* specific function, in departure from earlier reported redundancies. Furthermore, any differences in response to elf18 elicitation between WT and *tcp8* were abolished when leaf discs were pretreated with exogenous brassinolide (24-epiBL) (Figure 4B,C). Considering our findings that TCP8 binds and activates BR regulatory genes, these data suggest that this immune signaling phenotype is caused by a BR signaling deficiency in the *tcp8* background that is restored by the presence of excess BR and compensation by other signaling components.

**Figure 4.**
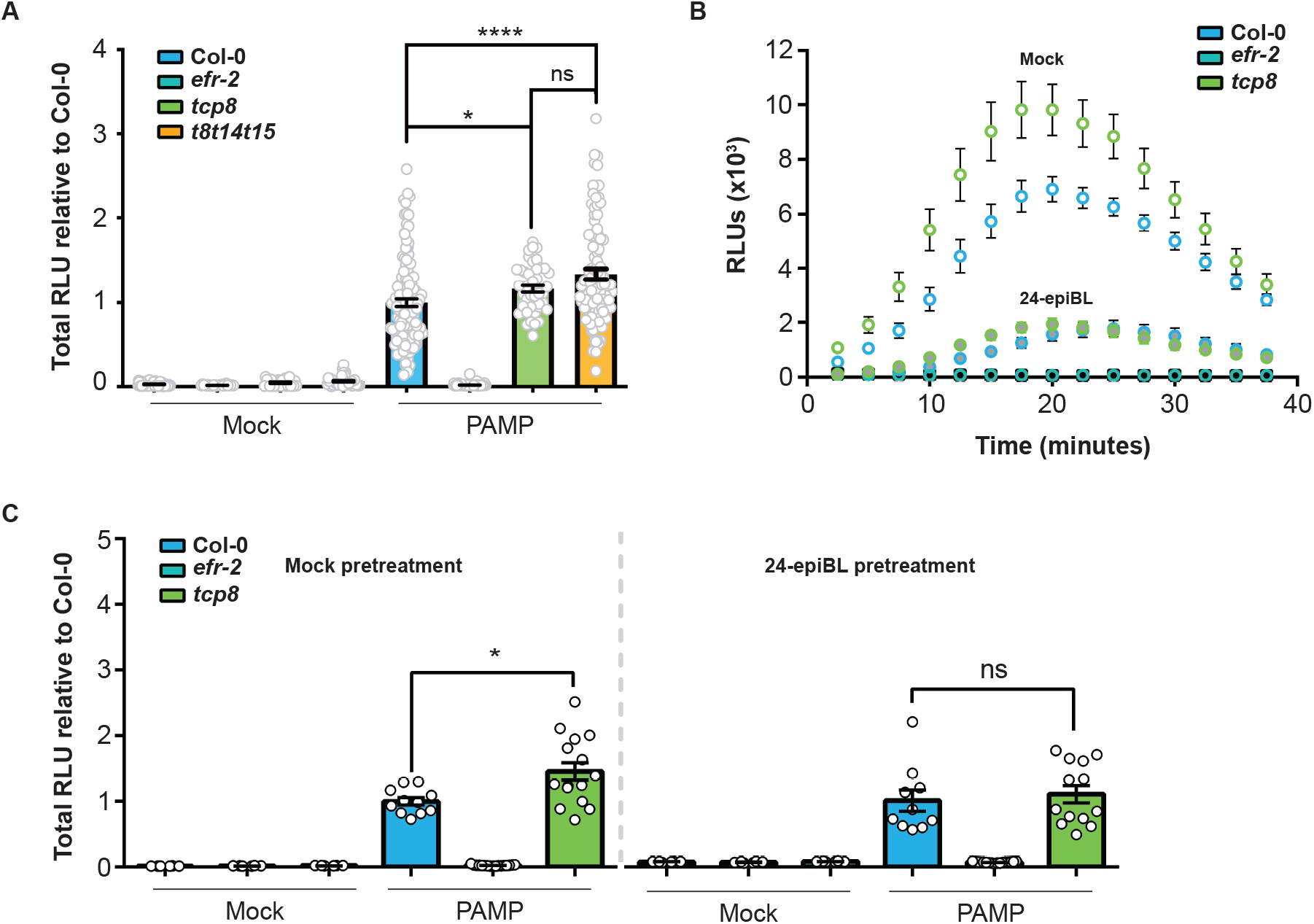
Enhanced elf18-induced ROS production in *tcp8* is suppressed by BR treatment. A, ROS production was elevated in *tcp8* relative to Col-0, but unchanged relative to *t8t14t15* after treatment with 1 μM elf18. Total RLUs were determined relative to Col-0 and data combined from 5 independent experiments, error bars indicate SE, \**P* <0.05, \*\*\*\**P* <0.0001 (n >46, Student’s *t*-test). B, In time course experiments, observed differences in ROS production between Col-0 and *tcp8* were attenuated by 24-hour pretreatment with 1 μM 24-epiBL *tcp8* (open shapes mock, filled shapes BL). Similar results were observed for 3 independent experiments. C, Total RLU measurements from (B) time course normalized to Col-0. \**P* <0.05 (*n* = 12-16, Student’s *t*-test).

### TCP8 regulates BR-responsive growth and directly interacts with BZR-family TFs

We reasoned that *tcp8* may exhibit signs of impaired BR signaling, such as altered BR-responsive growth patterns. However, no obvious morphological phenotypes were visible in the *tcp8* mutant, indicating that any insensitivity phenotypes were likely to be mild. In a standard BR-induced root growth inhibition assay in Arabidopsis, we found that *tcp8* seedlings exhibited relative root length (BL/mock) similar to a BR-insensitive *BZR2* rnai line and nearly twice the relative length of the fully sensitive Col-0 seedlings when grown on ½ MS plates containing 100 nM 24-epiBL (Figure 5A).

**Figure 5.**
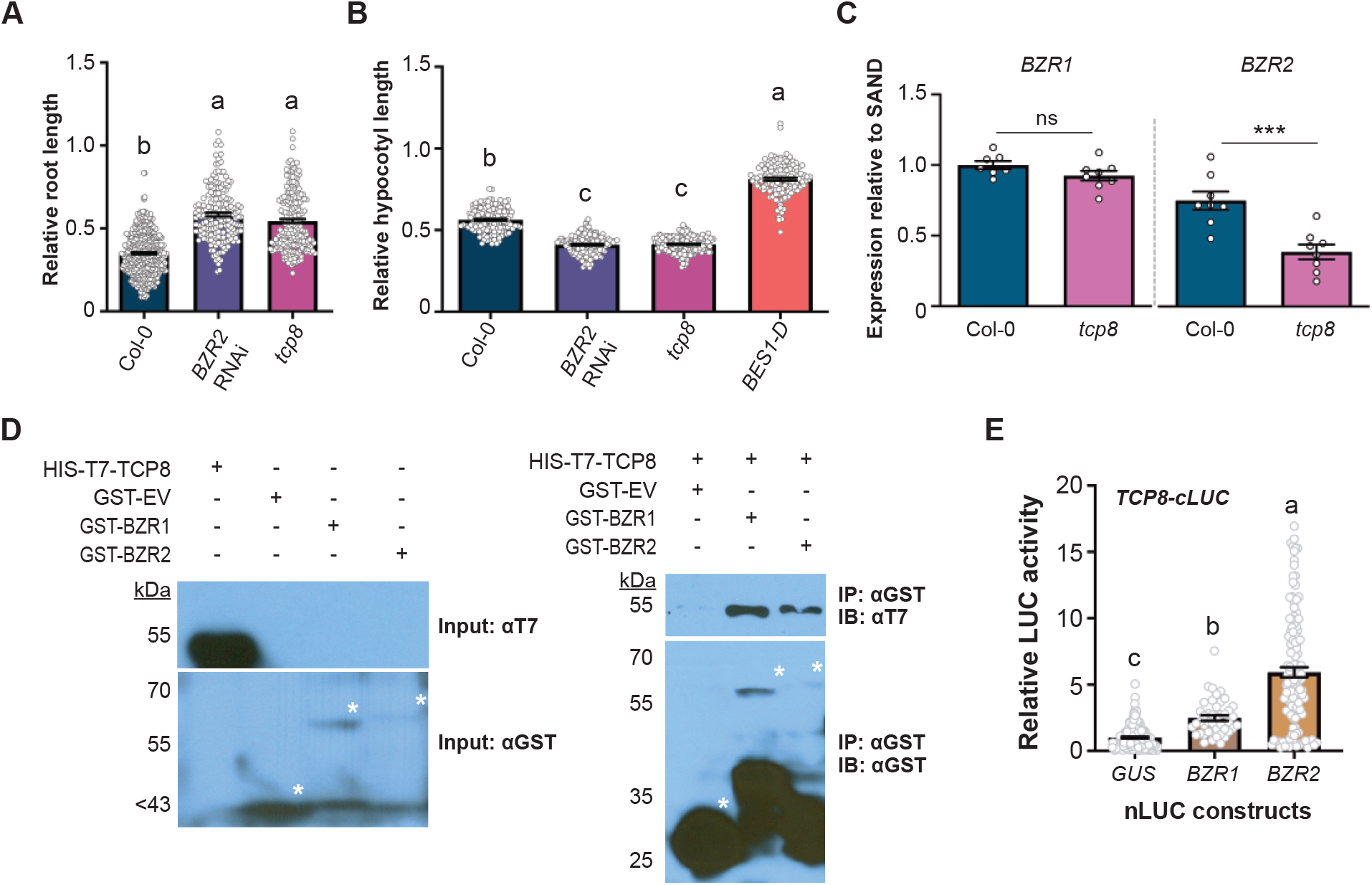
TCP8 controls BR-responsive growth patterns and interacts with BZR1 and BZR2. A, Seedlings were grown on ½ MS plates containing DMSO (mock) or 100 nM 24-epiBL in long-day growth conditions for 10 days. Proportional root length (BL treatment relative to mock) was determined for each genotype tested. Similar enhanced growth relative to Col-0 was observed in *tcp8* as in the BL-insensitive *BES1/BZR2* rnai line (*n* >184). B, Seedlings were grown on ½ MS plates containing DMSO or 500 nM Brz in darkness for 8 days. Proportional hypocotyl length (Brz treatment relative to mock) was determined for each genotype tested. Similar Brz hypersensitivity relative to Col-0 was observed in *tcp8* as in the *BES1/BZR2* rnai line, while the opposite effect was observed in dominant *bes1-D* mutant control line (*n* >134). For sensitivity experiments, bars indicate 1 SE, significance (ANOVA), *P* <0.01 with Tukey multiple comparison test. C, Transcript levels of *BZR2* are significantly reduced in *tcp8* relative to Col-0. Total RNA was collected from 10-day old seedlings grown under identical conditions as in (A). Expression levels normalized to the housekeeping gene *SAND* (*n* = 8). Error bars indicate SE, significance (ANOVA) with Tukey multiple comparison test, letters denote difference at *P* <0.001. D, *In vitro* interaction of HIS-T7-TCP8 with GST-BZR1 and GST-BZR2. The experiment was repeated twice with similar results. E, TCP8 interacts with BZR1 and BZR2 *in planta*. Interactions were evaluated by split-luciferase assay in *N. benthamiana* leaf epidermal cells. Total RLUs were normalized to non-interacting control (*nLUC-GUS*) levels as relative luciferase activity and data combined from 3 independent experiments. Error bars indicate 1 SE, significance (ANOVA) with Tukey multiple comparison test, letters denote difference at *P* <0.01 (*n* >130).

We hypothesized that impaired BR signaling in *tcp8* seedlings would result in predisposed sensitivity to inhibition of BR biosynthesis. Accordingly, we measured hypocotyl elongation of etiolated seedlings grown on ½ MS plates with 500 nM Brz. We observed a decrease in relative hypocotyl length (Brz/mock) in *tcp8* relative to fully sensitive Col-0 and the insensitive, constitutively active line *bes1-D*. Again, the *tcp8* phenotype mimics the hypersensitivity of the *BZR2* rnai line (Figure 5B).

To verify that loss of transcriptional activity in *tcp8* contributed to these phenotypes, we collected total mRNA from Arabidopsis seedlings grown on ½ MS under identical conditions to those in Figure 5A and measured the transcript levels of key TCP8-regulated BR genes. Although low basal *BZR1* expression levels likely dampened phenotypic differences, transcript levels were mildly reduced in *tcp8*, while *BZR2* transcripts were significantly reduced to nearly half of WT (Figure 5C). Furthermore, the full complementation of the 24-epiBL insensitivity phenotype by the *TCP8-HA* transgenic line suggests that the activity of TCP8 is sufficient for WT levels of responsiveness to BR (Supplemental Figure S3). Since *BZR1/BZR2* transcript levels are severely reduced in the *BZR2* rnai line, it is possible that the *tcp8* phenotype represents a combinatorial effect of mildly reduced transcription at several different BR signaling loci, or perhaps hints at a mechanism by which mild reduction of steady state expression may be phenotypically exacerbated by lack of signal amplification during stress.

The activities of brassinosteroid-responsive transcription factors are commonly regulated through interaction with other BR signaling proteins. As an example, *in vitro* and *in vivo* interactions with BZR1 and BZR2 have been demonstrated for the PIF family (Oh et al., 2012). TCPs heterodimerize and may form higher-order transcriptional regulatory complexes to regulate their activities; the co-occurrence of TCP binding sites with those of other TF families points towards this mechanism (Martin-Trillo and Cubas, 2010). The enrichment of G-box binding motifs in the immediate vicinity of our chIP-seq candidate peaks could suggest that TCP8 may bind those motifs directly. Another possibility is that TCP8 is capable of forming dynamic heteromultimeric regulatory complexes with other G-box binding TFs to regulate its gene targets, including BR-responsive genes.

To test this second mechanism, we cloned the coding regions of a subset of the BR-related genes identified in our chIP-seq analysis into GST-tagged *E. coli* expression vectors and performed pulldown assays with TCP8. Despite relatively low expression levels, GST-BZR1 and GST-BZR2 co-purified strongly with HIS-T7-TCP8 (Figure 5D).

Direct interactions were verified *in planta* using a split luciferase assay in *N. benthamiana* leaf epidermal cells. Using a C-terminally tagged TCP8-cLUC construct, strong interaction was observed with BZR2-nLUC and a weaker interaction with BZR1-nLUC relative to a non-interacting GUS-nLUC control (Figure 5E). The *in planta* results largely mirror the interactions observed in the *E. coli* system. These data point towards a model by which TCP8 directly promotes transcriptional activation of BR signaling pathways through direct and genetic interactions with master regulators *BZR1* and *BZR2*.

### TCP8 subnuclear localization and activities are BR-responsive

TCP8 and other class I TCP TFs have been observed to form nuclear condensates basally or as an induced response to various protein-protein interactions and stresses (Valsecchi et al., 2013; Kim et al., 2014; Mazur et al., 2017; Yang et al., 2017; Perez et al., 2019). The nature of these subnuclear localizations has yet to be explored, but could be sites of suppression, enhanced activation, or both at multiple genetic loci. Brassinosteroid-responsive TFs like the bHLH protein CESTA form similar nuclear condensates in a BR-dependent manner (Poppenberger et al., 2011). To determine the effect of brassinolide perception on the cellular localization of TCP8, *35S:GFP-TCP8* was transiently expressed in *N. benthamiana* leaf epidermal cells and the infiltrated tissue was treated with 2 μM Brz in order to reduce levels of endogenous BR signaling and gene expression. In Brz-treated cells, punctate localization of GFP-TCP8 was reduced relative to the mock treatment. However, elicitation of the Brz-treated cells with 1 μM 24-epiBL partially restored basal localization patterns, while a control treatment largely had no effect (Figure 6A). These data indicate BR perception may direct TCP8 localization into nuclear condensates and that endogenous levels of BR signaling may account for reports of similar TCP8 localization patterns without elicitation.

**Figure 6.**
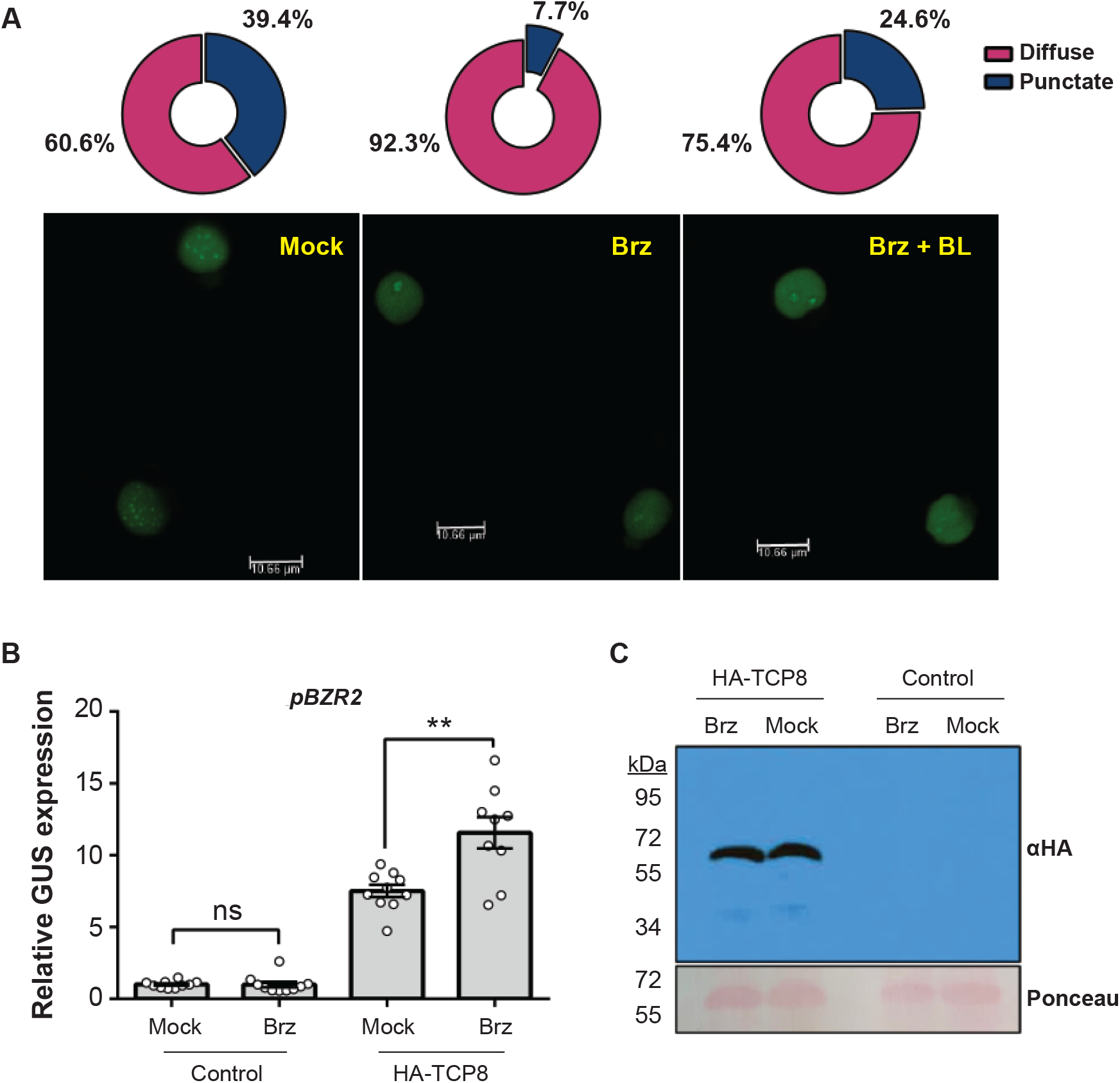
BR influences localization and activity of TCP in *N. benthamiana*. A, *35S:GFP-TCP8* was expressed in *N. benthamiana* leaf epidermal cells and leaves treated with either 2 μM Brz or DMSO mock solution, and then re-infiltrated with 1 μM 24-epiBL solution. Brz-treated samples exhibited significantly lower presence of nuclear condensates, while reapplication of 24-epiBL significantly increased these proportions, but not to the level of mock treatment (*n* > 69). *P* <0.0005 (Chi-Square Test for Association). B, Brz treatment enhances transactivation of the *pBZR2* promoter:GUS construct in *N. benthamiana*. GUS expression relative to control was measured after 24-hour pretreatment with 2 μM Brz or mock solution. Significantly higher GUS expression was observed in the Brz-treated leaf samples as compared to the mock treatment. Error bars indicate 1 SE, *\*\*P* <0.01 (*n* = 10, Student’s *t*-test). C, Total protein was isolated under the same conditions and HA-TCP8 protein levels analyzed by western blot with αHA antibody. Treatment with 2 μM Brz had no visible effects of TCP8 protein level relative to DMSO mock. Similar results were observed in 2 independent experiments.

In the case of CESTA, its BR-dependent movement into nuclear bodies is thought to suppress activation of BR gene expression (Khan et al., 2014). Along those lines, we explored the effects of Brz treatment on the transactivation by TCP8 in *N. benthamiana*. Interestingly, we observed increased activation of the *pBZR2* reporter construct by HA-TCP8 when leaves were treated with 2 μM Brz (Figure 6B). Total TCP8 abundance was unchanged by the treatment both in *N. benthamiana* (Figure 6C) and in the native promoter-driven Arabidopsis TCP8-HA line (Supplemental Figure S4), confirming that changes in relative activity were not a consequence of differences in protein level. Our data point towards a function for these observed nuclear condensates in the repression of TCP8 activities, demonstrated here in the context of BR signaling.

## DISCUSSION

The involvement of TCP8 and other class I TCPs in SA-dependent immune responses has been established in multiple recent studies, in part contributing to the family’s rise to prominence as a potential pathogen-targeted signaling hub. In one example, TCP14 is destabilized by the *P. syringae* (*Pst*) effector HopBB1 to antagonistically suppress SA and promote the virulence of the hemibiotrophic pathogen (Yang et al., 2017). More recently, TCP8 was demonstrated to be targeted by multiple effectors of another hemibiotroph, *X. campestris pv. campestris (Xcc*), but mechanisms of virulence have yet to be explored (González-Fuente et al., 2020). For any pathogen, virulence requirements extend beyond suppression of host immunity— this begs the question of what else these TCPs may be doing that make them such attractive targets. To fully understand the role of TCPs in these interactions, higher resolution of their transcriptional targets is required. We performed chIP-seq in this study with the goal of complementing our previous data supporting TCP8 as an effector target for its direct role in PTI. Several defense-genes were identified as TCP8 regulatory targets in our chIP-seq dataset, including WRKY-family TFs (*WRKY1, WRKY40*, and *WRKY51*), *ENHANCED DISEASE SUSCEPTIBILITY 1* (*EDS1*), and the NADPH oxidase *RbohF* (Table 1). Notably absent among the regulatory candidates is the receptor gene *EFR*, which is directly activated by TCP8 in response to elicitation by the PAMP elf18 (Spears et al., 2019). It is possible that untargeted chIP-seq may require elicitation to observe enrichment of that particular locus, or that we did not fully capture the scope of TCP8 binding targets with our use of a lower expression, native promoter-driven TCP8 transgenic line. This is perhaps reflected by the relatively low representation of bound defense gene promoters. A more complex model of gene regulation fits the limited overlap between our chIP-seq and RNA-seq data sets (a common observation). These binding sites may represent promoters that are bound but not constitutively activated or repressed, in wait of the appropriate stimulus, post-translational modification of TCP8, or activation of a necessary co-regulating TF partner as described for some WRKY TFs (Mao et al., 2011).

Interestingly, the list of regulatory candidates was dominated by hormone signaling pathway components, particularly those associated with auxin, gibberellin, and brassinosteroid signaling. This may point towards a role for TCP8 not as a primary regulator of any one particular phytohormone pathway, but rather as a regulator of baseline hormonal balancing across pathways; interactions with additional TF groups such as PIFs or BZRs may allow fine-tuning or specialization of TCP8 activity in response to environmental signals. This would support the proposed role of TCP8 and other class I TCPs as central signaling hubs, as well as ideal pathogen targets for manipulation of host hormone levels as a virulence mechanism.

In this manuscript, we focused primarily on the BR branch of regulatory candidates highlighted in the chIP-seq dataset, but the data sets generated will allow for additional studies to explore the involvement of TCP8 in other represented phytohormone pathways. Specifically, we describe a novel role of TCP8 as an activator of multiple key brassinosteroid master regulators, with activities and localization patterns controlled by cellular BR levels. These data highlight a TCP8-specific role that broadly contrasts with previous described functions. However, our data are in line with previous observations about its regulatory target BZR2 which is capable of promoting both BR-responsive gene expression and PTI-related gene expression, depending on the kinases it associates with and phosphorylation status at different residues (Kim et al., 2009; Kang et al., 2015). Mechanisms controlling its activation of multiple possible hormone-related signalling pathways are a key point of interest. One possibility is that TCP8 promoter occupancy switches between different hormone-responsive regulatory targets in response to local signals. Another could involve a core set of gene promoters that are involved in promoting multiple hormone signaling pathways themselves; a third mechanism previously described may include further regulation by combinatorial interactions between TCP8 and other TFs. These possibilities are by no means mutually exclusive.

It is possible that other TCP family or bHLH proteins may contribute to this phenotype by acting as a determining factor of which regulatory targets and pathways are activated by TCP8 under certain conditions. As with the regulation of skotomorphogenesis in the related PIF family (Zhang et al., 2013), TCP8 likely acts in concert with other TFs to cooperatively regulate a core set of BR genes at the level of hormone biosynthesis, signal transduction, and response. This is further supported by the finding that ∼25% of our identified TCP8-bound gene promoters (Supplemental Table S5) are also bound by BZR1 or BZR2 (Sun et al., 2010; Yu et al., 2011) and the observed *in vitro* and *in planta* interactions with BZR1 and BZR2 (Figure 5). Consistent with bHLH activities (Pireyre and Burow, 2015), combinatorial interactions with TF families regulating SA, auxin, or ABA signaling may contribute to the regulation of diverse biological processes by TCP8 that is reflected in our sequencing data. From the overabundance of auxin-responsive gene candidates identified in our chIP analysis (Figure 1), a role for TCP8 in auxin signaling would be particularly interesting in the context of this study due to known cooperation between auxin and BR and antagonism between auxin and SA that is thought to be exploited by pathogens as a virulence mechanism (McClerklin et al., 2018; Kong et al., 2020).

In the presence of BR, TCP8 may homodimerize or interact with TFs like BZR1 or BZR2 in heteromultimeric complexes to activate BR-responsive gene expression. Our data support a model by which increased concentrations of BR induce the movement of TCP8 into regulatory nuclear condensates to modulate BR biosynthesis or responsive gene expression. As BR levels are reduced, TCP8 may associate more weakly with repressive complexes, contributing to a generally diffuse localization pattern. In this state TCP8 can freely activate baseline subsets of BR genes, such as BZR2, to replenish hormone levels and activate other signaling responses induced under low-BR conditions (Figure 7). Similar instances of phase separation have already been characterized for transcriptional regulators of multiple phytohormone pathways (Powers et al., 2019; Zavaliev et al., 2020; Zhu et al., 2021). Although further study is required to clarify mechanisms and functions of TCP8 condensate formation, there is an intriguing possibility of similar induction by other TCP8-involved hormone signaling pathways. Additionally, as with the BR-dependent subnuclear localization of the bHLH CESTA (Khan et al., 2014), the formation of TCP8 condensates is likely regulated post-translationally. These findings highlight the potential significance of previously described TCP8 phosphosites (Xu et al., 2017) and sumoylation (Mazur et al., 2017) to its function in BR signaling. By this regulatory mechanism, TCP8 may act dynamically as one component of an elaborate ‘switch’ between different transcriptional priorities induced by a plant’s perception of defense or growth signals, or the yet uncharacterized manipulations of interacting pathogen effectors aiming to modulate phytohormone signaling outputs.

**Figure 7.**
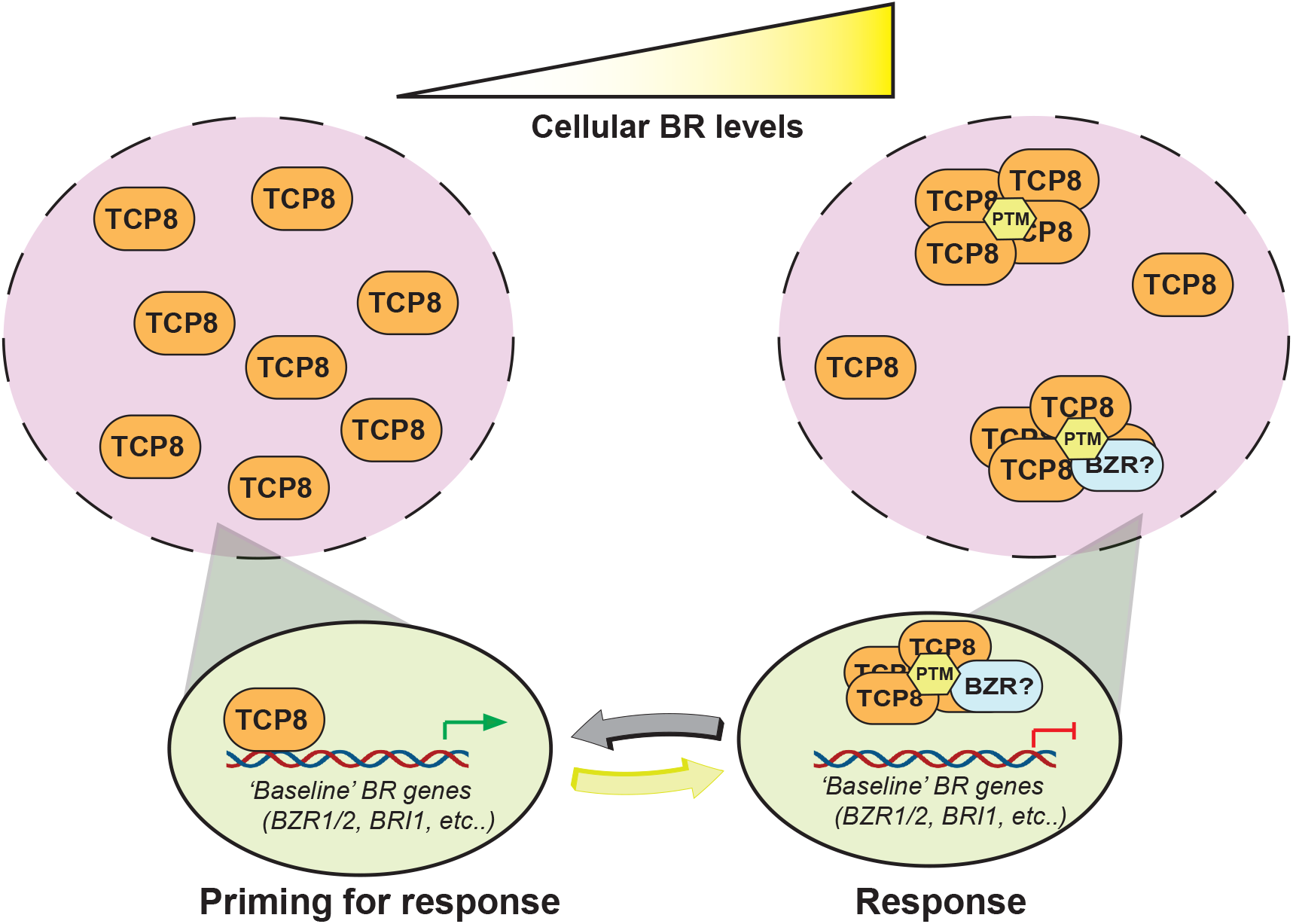
A working model for regulation of brassinosteroid signaling by TCP8 in Arabidopsis. At low levels of cellular BR, TCP8 associates with the promoters of ‘baseline’ BR genes to prime the cell for production of and response to BRs. At high levels of cellular BR, TCP8 involvement in the activation of BR signaling is no longer necessary, and excess TCP8 protein enters nuclear condensates, producing the ‘punctate’ nuclear phenotype observed in this study. Both the formation of these condensates and the described interactions with BR regulators BZR1 and BZR2 under basal conditions are likely dynamically regulated through post-translational modifications of TCP8. Although the function of these nuclear condensates remains to be explored, our data suggest they may be repressive in nature.

## MATERIALS AND METHODS

### Plant material and cultivation

Arabidopsis plants used in chIP-qPCR, chIP-seq, and RNA-seq experiments were grown in liquid half-strength Murashige Skoog (MS) media on a light rack under 24-hour light at room temperature. All other Arabidopsis plants were grown on soil or half-strength MS 0.8% phytoagar plates in an E-7/2 reach-in growth chamber (Controlled Environments Ltd.) under an 8-hour light and 16-hour dark cycle at 24°C, 70 to 80% relative humidity, and a light intensity of 140 to 180 μmol photons per square meter per second. *Nicotiana benthamiana* plants used in transactivation, interaction, and localization assays were grown under an 8-hour light and 16-hour dark cycle at 22°C and 55% relative humidity.

The transgenic *pTCP8:TCP8-HA* (Spears *et al*., 2018), *tcp8-1 and tcp* triple mutant (Kim et al., 2014), *efr-2* mutant (Zipfel et al., 2006), *BZR2* rnai (Yin et al., 2005) and *bes1-D* (Yin et al., 2002) lines have all been previously characterized.

### Molecular cloning

Full-length cDNAs of *BZR1* and *BZR2* without stop codons were amplified from cDNA template synthesized from Col-0 gDNA and cloned into the Gateway-compatible donor vector pDONR201. GUS and BR gene donor clones were moved by LR reaction into split luciferase vector pXnLUC, while a TCP8 donor clone was moved into pXcLUC.

BR gene donor clones were used as template for PCR-based addition of *BamHI* and *SmaI* restriction sites to the CDS. Products were digested and ligated into corresponding restriction sites in the GST-tag vector pGEX-4T3.

For transactivation experiments, 2 kb upstream regions of *BZR1* and *BZR2* were cloned into pDONR201. Mutant *tcp* variants of the promoters were generated by site-directed mutagenesis. Donor clones were moved by Gateway LR reaction into pYXT1 to produce complete promoter-reporter constructs.

### Assays in N. benthamiana expression system

For transactivation assays, *A. tumefaciens* strain GV3101 was electroporated with various promoter:GUS constructs, and strain C58C1 electroporated with a *35S:HA-TCP8* construct (Kim et al., 2014). Overnight cultures were generated for each transformant at 30°C, then pelleted and resuspended in 10 mM MgCl_2_ buffered with 1 mM MES (pH 5.6) and 100 nM acetosyringone (3’,5’-dimethyoxy-4’-hydroxyacetophenone). Suspensions were incubated for 4-5 hours. Bacterial inoculums were mixed at OD_600_ 0.2 for each strain and syringe infiltrated into mature *N. benthamiana* leaves. Tissue was collected 72 hours after inoculation with a 1 cm diameter hole punch, 4 leaf discs per sample. Protein was isolated in extraction buffer and GUS quantified according to a standard 4-MUG assay. Protein concentration was determined by Bradford assay and GUS levels calculated before normalization to the empty vector control. For Brz treatments, infiltrated leaves were then gently re-infiltrated with either a DMSO mock or 2 μM Brz solution, before incubating for an additional 24 hours.

Immunoblot analysis of epitope-tagged TCP8 in *N. benthamiana* and Arabidopsis was performed by extracting one gram of total protein in 1 mL of 2x sodium dodecyl sulfate (SDS) buffer (100 mM Tris-HCl, pH 6.8, 4% SDS, 20% glycerol, 250 mM dithiothreitol). Samples were cleared by centrifugation at 13,000 rpm and loaded onto an 8% bis-acrylamide SDS-polyacrylamide gel electrophoresis (PAGE) gel. Protein was detected with 1:2,000 horseradish peroxidase (HRP)-conjugated anti-HA antibody (Roche).

For BR localization assays, a *35S:GFP-TCP8* construct was transformed into *A. tumefaciens* C58C1 and inoculums prepared at OD_600_ 0.05. *N. benthamiana* leaves were syringe infiltrated and incubated for 48 hours under standard conditions. Infiltrated leaves were then gently re-infiltrated with either a DMSO mock or 2 μM Brz solution, before incubating for an additional 24 hours. 2 hours prior to observation under a Leica TCS SP8 microscope, leaves were infiltrated once more with either a DMSO mock or 1 μM 24-epiBL solution. Nuclei were randomly sampled (>20 per replicate) and GFP-TCP8 localization status evaluated for each. Effects of treatment on localization was determined by a Chi-Square Test of Association.

### Chromatin immunoprecipitation and chIP-seq analysis

Chromatin immunoprecipitation was performed with 10-day old seedlings grown in liquid ½ MS as previously described (Spears et al., 2019). For chIP-PCR, 5 μl of the resulting purified DNA was analyzed by qPCR as described above. For chIP-seq, chIP DNA and input DNA from 5 biological replicates were used to construct libraries with an NEBNEXT Ultra II Kit Library prep kit according to the manufacturer’s protocol. High-throughput sequencing of prepared libraries was performed on an Illumina NextSeq 500.

Processed FASTQ sequences were mapped to the Arabidopsis TAIR10 genome using Bowtie2. Non-uniquely mapped reads were removed, and enriched TCP8-binding peaks were called from pooled chIP/input files using MACS2 with estimated fragment size 189, shift size 0, and cutoff (FDR <0.05).

Genome distribution of called peaks was determined by the PAVIS tool (https://manticore.niehs.nih.gov/pavis2/) with upstream and downstream limits set to 3000 bp and 1000 bp, respectively.

Motif analysis of promoter-bound TCP8 binding peaks was performed using the MEME-Suite tools (https://meme-suite.org/meme/tools/meme). From upstream peak coordinates, 250 bp flanking sequences (500 bp total) were determined and analyzed using default settings.

Gene ontology (GO) term analysis was performed using the DAVID platform (https://david.ncifcrf.gov/).

### RNA sequencing and analysis

RNA was isolated from 10-day old seedlings grown under identical conditions as the chIP-seq experiments. Libraries were prepared using an Illumina TruSeq stranded library preparation kit according to manufacturer’s protocol. High-throughput sequencing of prepared libraries were performed on an Illumina NextSeq 500.

After sequencing adapters were removed, bases at the 5’ and 3’ end of each read were removed such that the confidence scores of each base in the remaining read were each above 99%. These trimmed reads were then mapped to the TAIR10 genome release using the ShortRead Package in R Bioconductor. Differential expression was assessed using the edgeR package, using tagwise dispersion estimates and Bonferroni and Hochberg multiple testing correction.

Ontology enrichment was performed using the GOStats package. The background frequency for each ontology (gene universe) was defined to be the union of all genes with (uncorrected) p-values ≤ 0.05, and all pairwise comparisons were made between genotypes.

### Quantitative real-time PCR

RNA was isolated from whole 10-day old seedlings under etiolating conditions and reverse transcription performed according to manufacturer specifications. qPCR was performed with 5 μl of 20x-diluted cDNA and a primer concentration of 1 μM in a 20 μl reaction with Brilliant III Ultra-Fast SYBR GREEN Master Mix (Agilent, www.agilent.com) with a BioRad CFX Connect thermocycler (Bio-Rad, www.bio-rad.com). Transcript levels were normalized to the housekeeping gene *SAND*.

### Arabidopsis BL/Brz sensitivity assays

BL sensitivity was evaluated by a standard root-length assay. Stratified seed was plated on square ½ MS plates with either mock DMSO or 100 nM 24-epiBL (*Sigma E1641*). Plates were placed vertically at 4 °C for two days with foil covering before removal of foil and transfer to a short-day growth chamber. Seedlings were grown for approximately 10 days before root length was measured using ImageJ.

Brz sensitivity was evaluated by a similar protocol to measure etiolated hypocotyl elongation. Briefly, stratified seed was plated on square ½ MS plates with either mock DMSO or 500 nM Brz (*Sigma SML1406*). Plates were covered in foil and placed vertically at 4 °C for two days before removing the foil and transferring plates to a short-day growth chamber for 3 hours. Plates were then covered again in foil and seedlings were grown for 8 days before hypocotyl length was measured using ImageJ.

### Arabidopsis immunity assays

ROS production was measured by a standard H_2_O_2_-dependent luminescence assay (Heese et al., 2007) with modifications. Briefly, 1 cm diameter Arabidopsis leaf discs were halved and then floated in 100 μl water with either DMSO or mock or 1 μM 24-epiBL added in a 96-well plate under full light overnight. Incubation solution was removed and replaced with 100 nM elf18 or DMSO mock elicitation solution. Luminescence was quantified in 2-minute increments over a 30-minute period immediately after elicitation.

Bacterial growth assays were performed as previously described (Spears et al., 2019).

### Protein interaction assays

*In vitro* co-pulldown assays with GST-EV/BZR1/BZR2 and His-T7-TCP8 were performed as previously described (Halane et al., 2018). Briefly, GST-tagged proteins were pulled down, incubated with HIS-T7-TCP8 lysate, and eluted from binding columns before probing with αT7 and αGST antibodies.

*In planta* split luciferase assays were performed according to a standard protocol (Cazzonelli and Velten, 2006) with modifications. Briefly, GUS/BZR1/BZR2-nLUC and TCP8-cLUC constructs were transformed into *A. tumefaciens* C58C1 and co-inoculated at OD_600_ 0.2 into *N. benthamiana* leaves by syringe infiltration. After 72 hours, leaf discs were taken by a sharp 0.5 cm diameter (#2) bore and floated abaxial side down on 100 μl infiltration solution (50 mM MES pH 5.6, 10 mM MgCl_2_, 0.5% DMSO) in a white 96 well plate with lid. Plates were wrapped in foil and incubated in a growth chamber for 20 minutes. Infiltration solution was removed by multipipettor and replaced with 100 μl reaction solution (1x infiltration solution, 1mM luciferin (Goldbio #LUCK-100)). Luminescence was quantified in 10-minute increments over a 2-hour period in a BioTek Synergy HTX plate reader.

### Accession Numbers

The TAIR accession numbers for referenced genes are as follows:

At1g09530 (*PIF3*), At1g19350 (*BZR2*), At1g25330 (*CESTA*), At1g31880 (*BRX*), At1g58100 (*TCP8*), At1g69690 (*TCP15*), At1g73830 (*BEE3*), At1g74710 (*ICS1*), At1g75040 (*PR5*), At1g75080 (*BZR1*), At1g80840 (*WRKY40*), At2g28390 (*SAND*), At3g47620 (*TCP14*), At4g16890 (*SNC1*), At4g39400 (*BRI1*), At5g20480 (*EFR*), At5g46330 (*FLS2*).

## Data Availability

The sequencing data used in this study are openly available in the NCBI SRA at https://www.ncbi.nlm.nih.gov/sra, accession number PRJNA754790.

## Acknowledgements

We are grateful to L. Cseke for assistance in manuscript review and preparation, to C. Garner for technical advice and microscopy assistance and to P. Martin and M. Creach for plant maintenance and general assistance. Thanks to the University of Missouri Molecular Cytology Core for technical assistance and setup involved in acquiring confocal images presented in this study. We also thank M.Y. Zhou and the University of Missouri Genomics Technology Core for technical support and services involved in Illumina library preparation and sequencing.

## Supporting Information

**Figure S1** Venn diagrams of class I TCP regulatory target groups.

**Figure S2** Bacterial growth levels are unaffected in *tcp8* relative to Col-0.

**Figure S3** Affected BR-responsive growth patterns are restored in the *TCP8-HA* complemented line.

**Figure S4** TCP8 protein levels are unaffected by cellular BR levels in Arabidopsis.

**Table S1** TCP8 binding peaks identified by chIP-sequencing.

**Table S2** TCP8 binding peaks categorized by genomic location.

**Table S3** DEGs identified by RNA-sequencing in *tcp8* and *t8t14t15*.

**Table S4** TCP8-bound DEGs identified by RNA-sequencing in *tcp8* and *t8t14t15*.

**Table S5** Comparison between known TCP8 and BZR gene promoter targets

**Table S6** Primers used in this study

